# Oxygen Restriction Generates Difficult-To-Culture Pathogens

**DOI:** 10.1101/339747

**Authors:** Lasse Kvich, Blaine Fritz, Stephanie Crone, Kasper N. Kragh, Mette Kolpen, Majken Sønderholm, Mikael Andersson, Anders Koch, Peter Ø. Jensen, Thomas Bjarnsholt

**Author notes:** **Correspondence**: Professor Thomas Bjarnsholt, Costerton Biofilm Center, Department of Immunology and Microbiology, University of Copenhagen, Blegdamsvej 3b, Room 22.2.22, DK-2200 Copenhagen N, Denmark., Associated Professor Peter Ø. Jensen, Costerton Biofilm Center, Department of Immunology and Microbiology, University of Copenhagen, Blegdamsvej 3b, Room 24.1.14, DK-2200 Copenhagen N, Denmark.

## Abstract

Induction of a non-culturable state has been demonstrated for many bacteria. In a clinical perspective, the lack of growth due to these non-culturable bacteria can have major consequences for the diagnosis and treatment of patients. Here we show how anoxic conditioning (restriction of molecular oxygen, O_2_) generates difficult-to-culture (DTC) bacteria during biofilm growth. A significant subpopulation of *Pseudomonas aeruginosa* entered a DTC state after anoxic conditioning, ranging from five to 90 % of the total culturable population, in both planktonic and biofilm models. Anoxic conditioning also generated DTC subpopulations of *Staphylococcus aureus* and *Staphylococcus epidermidis* (89 and 42 % of the total culturable population, respectively). Growth of the DTC populations were achieved by substituting O_2_ with 10 mM NO_3_^-^ as an alternative electron acceptor for anaerobic respiration or, in the case of *P. aeruginosa*, by adding sodium pyruvate or catalase as scavengers against reactive oxygen species (ROS) during aerobic respiration. An increase in normoxic plating due to addition of catalase suggests the molecule hydrogen peroxide as a possible mechanism for induction of DTC *P. aeruginosa.* Anoxic conditioning also generated a true viable but non-culturable (VBNC) population of *P. aeruginosa* that was not resurrected by substituting O_2_ with NO_3_^-^ during anaerobic respiration. These results demonstrate that habituation to an anoxic micro-environment could complicate diagnostic culturing of bacteria, especially in the case of chronic infections where oxygen is restricted due to the host immune response.

**Importance:** Diagnostics of bacteria from chronic infections by standard culture-based methods is challenging. Bacteria in a non-culturable state may contribute to the lack of culturing from these infections. Many stressors are known to induce a non-culturable state, among others the absence of molecular oxygen, which is evident in chronic infections due to high rates of oxygen consumption by the host response. In this study, we have shown that *Pseudomonas aeruginosa, Staphylococcus aureus* and *Staphylococcus epidermidis* can enter a non-culturable state after oxygen restriction. Regrowth was not possible using conventional normoxic plating where oxygen served as electron acceptor. Instead, regrowth was enabled during anoxic conditions with added nitrate as alternative electron acceptor. In the case of *P. aeruginosa* we were able to counteract this non-culturable state by adding scavengers for reactive oxygen species during normoxic incubation. This suggest a link between reactive oxygen species and a non-culturable state and it also renders a solution to circumvent this problem in a clinical diagnostic setting. Our findings show that bacteria can habituate to their environment and that it has to be taken into consideration especially when culturing clinical samples e.g. from chronic infections.

## Introduction

Clinical laboratories often culture bacteria with enriched media, developed to support maximal growth of particular pathogens. However, even these bacterial human pathogens can enter a growth-restricted, transient state resulting in loss of culturability, especially following antibiotic treatment (Pasquaroli et al., 2013). Non-culturable bacteria are apparent by microscopic identification in clinical cases presenting sign of infection, but no positive cultures. These cases are especially prevalent during chronic infection, where lack of culturability prevents proper diagnosis and treatment (Høiby et al., 2010;Costerton et al., 2011;Høiby et al., 2015).

Two well defined, yet difficult to distinguish, non-culturable states have been described for bacteria: viable but non-culturable (VBNC) and persisters (Oliver, 2010;Wood et al., 2013;Fisher et al., 2017;Kim et al., 2018). Several environmental stresses have been associated with non-culturable states, e.g. oxidative stress, change in temperature or pH, nutrient starvation, change in osmotic concentrations and presence of heavy metals or antibiotics (Oliver, 2010;Ayrapetyan et al., 2015). Oxygen starvation, referred to as anoxic conditioning in this paper, has also shown to induce a non-culturable state in batch cultures of *Pseudomonas aeruginosa*, but supplementation with nitrate (NO_3_^-^) as an alternative electron acceptor during anoxic plating restored culturability (Binnerup, 1993). The lack of growth during normoxic plating suggests that anoxic conditioning sensitizes a subpopulation of *P. aeruginosa* to O_2_ or its toxic derivatives, such as reactive oxygen species (ROS). ROS are continuously created during aerobic respiration by incomplete reduction of O_2,_ which is toxic to cells if not scavenged (Fenchel and Finlay, 2008). Presence of ROS has, in some studies, been shown to generate VBNC bacteria (Oliver, 2010;Noor, 2015), but most research has focused on *Escherichia coli* strains or *Vibrio* spp., while the effects of ROS on *P. aeruginosa* and other facultative pathogens are not well characterized.

Evidence of anoxic zones and anaerobic bacterial activity in chronic infections suggests that colonizing bacteria, such as *P. aeruginosa*, experience anoxic conditioning (Hassett et al., 2002;Worlitzsch et al., 2002;Kolpen et al., 2014). High rates of O_2_ consumption by polymorphonuclear leukocytes generate local, anoxic zones and likely play a major role in O_2_ depletion during chronic infection (Høiby et al., 2015). In addition, other cells of the host may also consume O_2_, increasing the likelihood of bacteria experiencing anoxia (Worlitzsch et al., 2002). Many chronic infections are thought to contain bacteria in the biofilm mode of growth (Costerton et al., 2003) and endogenous O_2_ depletion inside the biofilm (Sønderholm et al., 2018), along with intense O_2_ consumption by the host immune response (Worlitzsch et al., 2002;Kolpen et al., 2010;Trunk et al., 2010;Kragh et al., 2014;James et al., 2016;Jensen et al., 2017), may therefore increase the number of VBNC bacteria.

This study aimed to determine whether anoxic conditioning generates VBNC cells of *P. aeruginosa* during biofilm growth. Furthermore, we investigated whether ROS are involved in the loss of culturability.

## Materials and Method

### Bacterial strains

*Pseudomonas aeruginosa* (PAO1, ATCC 15692), a catalase A deficient *Pseudomonas aeruginosa* strain (Δ*katA* PAO1) (Hassett et al., 1999), *Staphylococcus epidermidis* ATCC 14990, *Staphylococcus aureus* NCTC 8325-4 (methicillin susceptible) (Frees et al., 2004), *Staphylococcus aureus* USA300 JE2 (MRSA) (Fey et al., 2013), E*scherichia coli* CFT073 (Welch et al., 2002) and a clinical strain of *Enterococcus faecalis* from the Department of Clinical Microbiology, Copenhagen University Hospital – Rigshospitalet, Denmark were used in this study.

### Agar plates and media

Primarily lysogeny broth (LB) agar plates were applied in this study. Lysogeny broth (pH 7.5) consisted of 5 g/L yeast extract (Oxoid, Roskilde, Denmark), 10 g/L tryptone (Oxoid), 10 g/L NaCl (Merck, USA). All plates contained 2 % agarose. Plates used for anoxic growth were supplemented with 10 mM KNO_3_ (Sigma-Aldrich, USA) to serve as alternative electron acceptor during anoxic plating, referred to as NO_3_^-^ throughout the paper. All agar plates and media in this study were supplied by the Panum Institute Substrate Department (Copenhagen, DK).

### Anoxic growth

Experiments investigating growth under anoxic conditions were performed in an anaerobic chamber (Concept 400 Anaerobic Workstation, Ruskinn Technology Ltd, UK). The gas atmosphere consisted of N_2_/H_2_/CO_2_ (ratio - 80:10:10). Anoxic chamber environment was confirmed with an oxygen sensor (HQ40d multi, HACH Company, USA). All media and chemical solutions used in anaerobic experiments were equilibrated in the anaerobic chamber 3 days prior to experiment. In the case of solutions requiring refrigeration, a minimal volume was applied (< 1 mL), sealed with Parafilm M, and thoroughly shaken upon entry into anaerobic chamber for quick gas equilibration.

### Direct viable count with LIVE/DEAD staining

LIVE/DEAD staining was applied to estimate the proportion of viable and non-viable cells in 24-hour-old batch cultures and in 16-day-old anoxically conditioned batch cultures of *P. aeruginosa*. The dyes consisted of two fluorescent nucleic acid stains, the green fluorescent stain (live cells) SYTO9 (Invitrogen, USA) and the red fluorescent stain (dead cells) propidium iodide (PI, Sigma-Aldrich, USA). SYTO9 penetrates both intact and damaged membranes while PI only stains damaged cells, thereby creating an opportunity to discriminate between live and dead cells (Li et al., 2014). Bacterial suspensions were vortexed thoroughly (1 min) and sonicated (5 min degas + 5 minutes sonication) before staining. To stain the cells, one µL of PI and SYTO9 was added to 1 mL of bacterial suspension and incubated for 15 minutes at room temperature. Suspensions of each biological replicate were then filtered through a 0.2 µm black Whatman, Nuclepore Trach-Etch Membrane (Sigma-Aldrich, Denmark) and visualized with confocal laser scanning microscopy using a Zeiss LSM 710 with a 63×/1.4 (numerical aperture) objective (Zeiss, Germany). Fifteen random fields (135µm × 135 µm) were examined for each filter (n=3). Enumeration of live (green) and dead (red) bacteria were done with the IMARIS software package (Bitplane AG, Schwitzerland). Briefly, individual particles in each field were identified and counted as bacteria. Particles that were stained with both PI and SYTO9 were considered non-viable and thus counted as dead cells. Conversion of enumerated viable and dead cells to bacterial counts per milliliter was performed as described (Boulos et al., 1999) to compare them with CFU/mL. Briefly, the numbers of particles per mL were estimated using the following equation 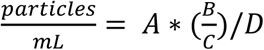. Where A is the amount of particles per field, B is the surface of filtration (mm), C is the area of the microscopic field and D the volume of the sample filtered. Normoxic and anoxic CFU/mL was carried out simultaneously to estimate the proportion of VBNC cells. CFU/mL was determined in the same way as described for liquid batch cultures.

### Bead-embedded inoculum of *P. aeruginosa*

Preparation of alginate beads with *P. aeruginosa* was carried out according to a method described by Sønderholm *et al*. (Sønderholm et al., 2017). Subsequently, beads were divided (10 beads per vial) into vials (Oximate Vial, PerkinElmer Inc., USA) containing 15 mL LB medium. Vials were sealed with Parafilm M (oxygen permeable) and incubated at 37^°^C on an orbital shaker at 180 rpm. Anoxic and normoxic CFU counts were determined from beads and from the surrounding suspension from the same vials on each sampling day (day 3, 6, 8, 12, 19 and 21). Two beads were sampled per biological replicate (n=4). Before determination of CFU, the beads were washed twice with 0.9 % NaCl to remove non-attached cells and transferred to 1.5 mL microcentrifuge tubes (Sigma-Aldrich, USA). One-hundred microliters of 0.1 M sodium carbonate (Na_2_CO_3_) followed by 100 µL of 0.04 M citric acid was added to the tubes to dissolve the beads. The suspensions were then sonicated (5 min degas + 5 minutes sonication; Bransonic ultrasonic cleaner 2510, Emerson Electric, USA) before ten-fold dilution series were performed in 0.9 % NaCl. CFU was determined by plating three, 10 µL-drops per dilution per replicate. Anoxic CFU determinations were performed inside an anaerobic chamber on LB plates supplemented with 10 mM NO_3_^-^. The same dilution series were applied to normoxic CFU determination outside the anaerobic chamber on LB plates with 10 mM NO_3_^-^. Plates were incubated 2 days before counting CFU.

### Filter biofilms with *P. aeruginosa*

This protocol was adapted to grow reproducible biofilms under anoxic and normoxic conditions. The method has previously been described by Bjarnsholt *et al.* (2015). The filter biofilms were kept on the same LB plates throughout the experiment. *P. aeruginosa* was propagated from frozen stock and grown overnight in 20 mL LB medium at 37^°^C on an orbital shaker at180 rpm. Cultures were adjusted to an optical density of 0.05 (OD_600_; UV spectrophotometer UV-1800 UV-VIS, Shimadzu corporation, JP) and 10 µL was transferred to the cellulose nitrate membrane filters (25 mm in diameter, GE Healthcare Life Sciences, UK). Plates were incubated under normoxic and anoxic conditions and kept in plastic bags with wet paper to avoid dehydration. Two filters were sampled per biological replicate (n=4) on each sampling day (day 1, 3, 7, 15 and 17). Filters were removed, placed in 10 mL tubes containing 5 mL 0.9 % NaCl, vortexed thoroughly (one min.) and sonicated (5 min degas + 5 minutes sonication). CFU/mL was determined as previously described.

### Liquid batch cultures of *P. aeruginosa*

*P. aeruginosa* was propagated from frozen stock and grown overnight in 20 mL LB media at 37°C and orbitally shaken at 180 rpm. Cultures were adjusted to an optical density (OD_600_) of 0.1 in glass vials (Oximate Vial) with a final volume of 20 mL LB. Vials were left to incubate at 37°C and orbitally shaken at 180 rpm. To create a normoxic environment, half of the vials were incubated with Parafilm M on top (normoxic conditioning), while the rest were incubated with a lid on top creating an anoxic environment (anoxic conditioning). Anoxic and normoxic plating was carried out from normoxic and anoxic conditioned liquid batch cultures (referred to as batch cultures throughout the paper) on each sampling day (day 1, 3, 5, 9, 11, 14, 16, 18, 21 and 28). Normoxic plating was carried out on LB plates with and without 10 mM NO_3_^-^ to test whether presence of NO_3_^-^ affected the number of CFU. Two mL (2 × 1 mL = 2 technical replicates) were sampled per biological replicate (n=4) on each sampling day. CFU/mL was determined as previously described.

### Colonies of *P. aeruginosa*

*P. aeruginosa* was propagated from frozen stock and grown overnight in 20 mL LB media at 37°C on an orbital shaker at 180 rpm. The cultures were then streaked onto LB plates supplemented with 10 mM NO_3_^-^. Plates were incubated under normoxic and anoxic conditions and kept in plastic bags with wet paper to avoid dehydration. A one-μl loop was used to sample colony material from each biological replicate (n=3) on each sampling day (day 6, 9, 13, 15 and 20). Colonies were transferred to 1.5mL microcentrifuge tubes (Sigma-Aldrich, Denmark) with 0.5 mL 0.9 % NaCl, vortexed thoroughly (1 min) and sonicated (5 min degas + 5 minutes sonication). CFU/mL was determined as previously described.

### Reactive oxygen species (ROS)

To elucidate whether growth of anoxically conditioned bacteria was restricted by creation of ROS during aerobic respiration, sodium pyruvate and catalase (Sigma-Aldrich, USA) was tested as ROS scavenger in LB plates with 10 mM NO_3_^-^. Sodium pyruvate (0.3 %) was tested during anoxic and normoxic plating for 24-hour-old batch cultures and anoxically conditioned 16-day-old batch cultures of *P. aeruginosa*. Catalase (∼50,000 units of catalase per L medium) was tested during anoxic and normoxic plating for anoxically conditioned 16-day-old batch cultures of *P. aeruginosa*. Furthermore, CFU/mL was also determined from 16-day-old anoxically conditioned batch cultures of *P. aeruginosa* and a catalase A deficient *P. aeruginosa* (Δ*katA* PAO1) to test the influence of ROS. Lastly, creation of ROS was measured along with optical density (OD) at day 1, 8 and 16 for normoxically and anoxically conditioned batch cultures of *P. aeruginosa*. Creation of ROS was measured by adding 5 µM 2′,7′dichlorodihydrofluorescein diacetate (DCHF-DA; Sigma) to the bacterial suspension. This concentration has previously been used to detect ROS for *P. aeruginosa* (Jensen et al., 2014). Two hundred microliter of anoxically conditioned cultures (OD_600_ adjusted to 0.1 in fresh LB medium) was added to each well of black 96-well microtiter plates with transparent flat bottom (16503, Thermo Fisher Scientific, Rochester, NY). OD and ROS was measured over 15 hours at 27°C with a VICTOR Multilabel Plate Reader (Perkin Elmer, MA). All experiments consisted of 3-4 biological replicates.

### Filter biofilms with other pathogens

To determine if other bacteria behaved in the same way as *P. aeruginosa*, we investigated the effect of anoxic conditioning on a selected group of pathogens (*Staphylococcus epidermidis* ATCC 14990, *Staphylococcus aureus* NCTC 8325-4 (methicillin susceptible), *Staphylococcus aureus* USA300 JE2 (MRSA), E*scherichia coli* CFT073 and a clinical strain of *Enterococcus faecalis*). The filter biofilm method was used as described earlier. For the methicillin susceptible strain, CFU was determined in the same way as described for the filters with *P. aeruginosa*. For the remaining strains, CFU was determined at day 1 and 9 on LB plates supplemented with 10 mM NO_3_^-^ ± O_2_. Furthermore, CFU was determined on LB plates supplemented with 10 mM NO_3_^-^ and 0.3 % sodium pyruvate to evaluate the effect of ROS. A minimum of three biological replicates was performed and CFU was carried out in the same way as described earlier.

### Statistics

To evaluate the difference in bacterial growth between incubation conditions (normoxic and anoxic conditioning), a linear regression was used with the difference between the logarithmically transformed values for normoxic and anoxic plating as outcome and with day (categorical) and interaction between day and plating condition (binary) as explanatory variables in SAS Genmod Procedure. The p-value of the interaction term was used as the p-value for the difference in bacterial growth. Log difference was calculated as (log_10_[CFU/mL]_anoxic_ - log_10_[CFU/mL]_normoxic_). The mean and standard error of the mean (SEM) were calculated for recovering bacteria and plotted using GraphPad Prism 6.1 (GraphPad Software, La Jolla, USA). From the ratio between anoxic (CFU - O_2_) and normoxic (CFU +O_2_) colony counts it was possible to calculate the fraction of DTC bacteria 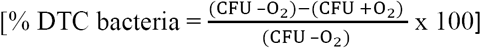. Data that were not part of the long-term experiments were instead analyzed with a 1-way ANOVA followed by Tukey’s multiple comparison tests or an unpaired t-test. A p-value ≤ 0.05 was considered statistically significant. The tests were performed with either Prism 6.1 (GraphPad Software, La Jolla, USA) or SAS v.9.4 (SAS Institute Inc., Cary, NC, USA).

## Results

### Direct viable counts with LIVE/DEAD staining revealed a VBNC *P. aeruginosa* population

Direct viable counts were carried out for 16-day-old, anoxically conditioned batch cultures of *P. aeruginosa* to investigate if anoxic conditioning could generate a VBNC population. Findings were compared to normoxic and anoxic plating (plating methods) performed on LB plates supplemented with 10 mM NO_3_^-^ (Fig. 1B). Anoxic plating yielded a significantly (p = 0.023) higher count of colony forming units per milliliter (CFU/mL) than normoxic plating (0.85 log values ± 0.04 SD). When comparing bacterial plate counts with direct viable counting, significantly more viable cells were measured compared to normoxic (2.31 log values ± 0.47 SD, p < 0.0001) and anoxic plating (1.46 log values ± 0.5 SD, p = 0.0009). The cells represented by this difference were considered VBNC. Direct viable counts were also performed on 24-hour-old batch cultures to investigate whether findings of VBNC *P. aeruginosa* were restricted to anoxically conditioned cells. No difference in bacterial counts was observed (Fig. 1A).

**Fig. 1.**
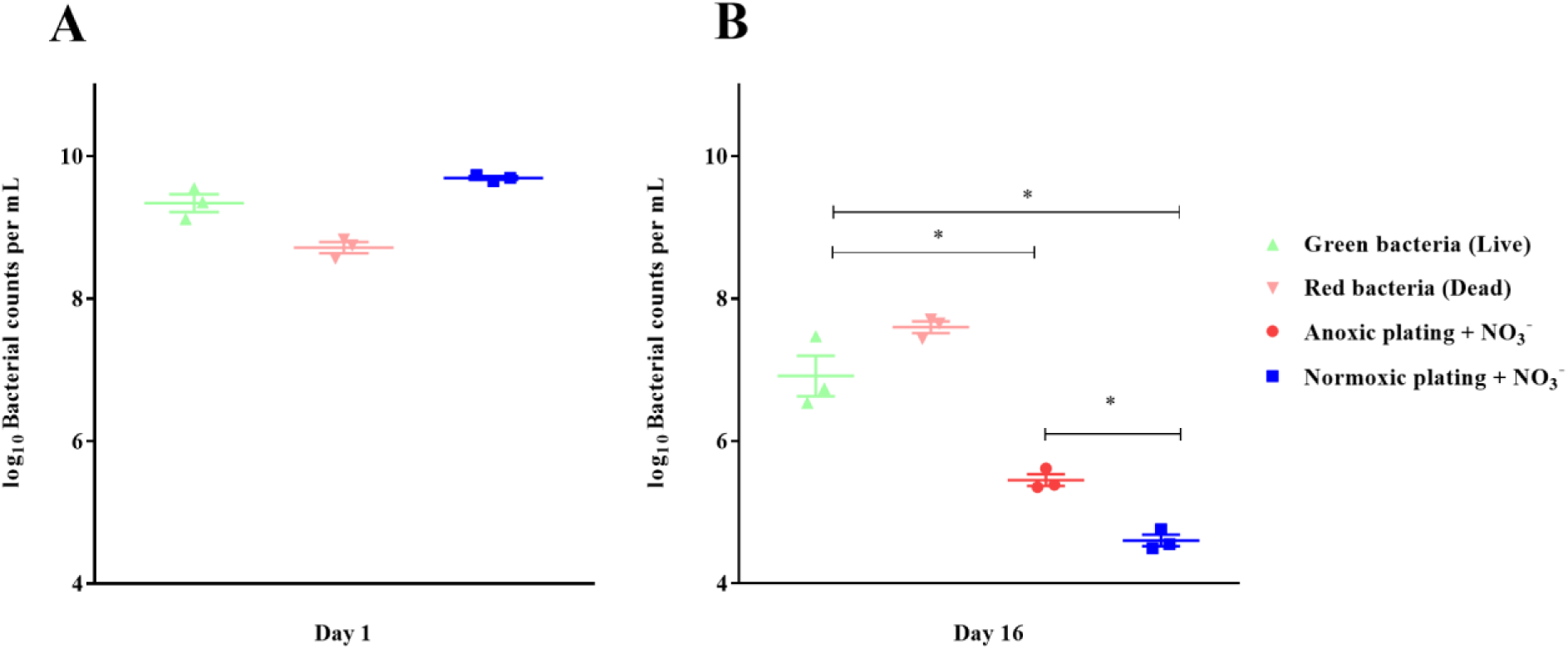
Direct viable counting reveals a viable but non-culturable population of *Pseudomonas aeruginosa* (PAO1) after anoxic conditioning. Bacterial counts per milliliter were determined with plate counting and direct viable counting from 24-hour-old batch cultures (A) and from 16-day-old anoxic conditioned batch cultures (B) of PAO1. Bacteria were stained with LIVE/DEAD staining to estimate the proportion of viable (“live”) and non-viable (“dead”) cells. There was significantly more viable bacterial counts when applying direct viable counting in comparison to normoxic plating (p < 0.0001, one-way ANOVA test) and anoxic plating (p = 0.0009, one-way ANOVA). Anoxic plating yielded significantly higher bacterial counts (p = 0.023, one-way ANOVA test) than normoxic plating. Symbols with error bars indicate the mean ± SEM (n = 3). + NO_3_^-^ refers to the addition of 10 mM KNO_3_ to LB agar plates.

### Anoxic conditioning generated oxygen intolerant subpopulations of *P. aeruginosa* in biofilm and planktonic models

To examine if biofilm growth affected the subsequent normoxic and anoxic plating, *P. aeruginosa* was grown in an alginate-bead biofilm model (Sønderholm et al., 2017). Plate counts were performed on LB plates supplemented with 10 mM NO_3_^-^. Plates were incubated under normoxic and anoxic conditions and plate counts were compared from the beads (biofilm) and the surrounding media (planktonic) of the same vials over a period of 21 days (Fig. 2A and 2B, respectively). The log difference was significantly (p = 0.003) higher for biofilm than planktonic cells (Fig. 2C). The cells represented by this difference were considered to be in a difficult-to-culture (DTC) state and ranged from five to 54 % of the entire population from day six to 19 (Fig. S1).

**Fig. 2.**
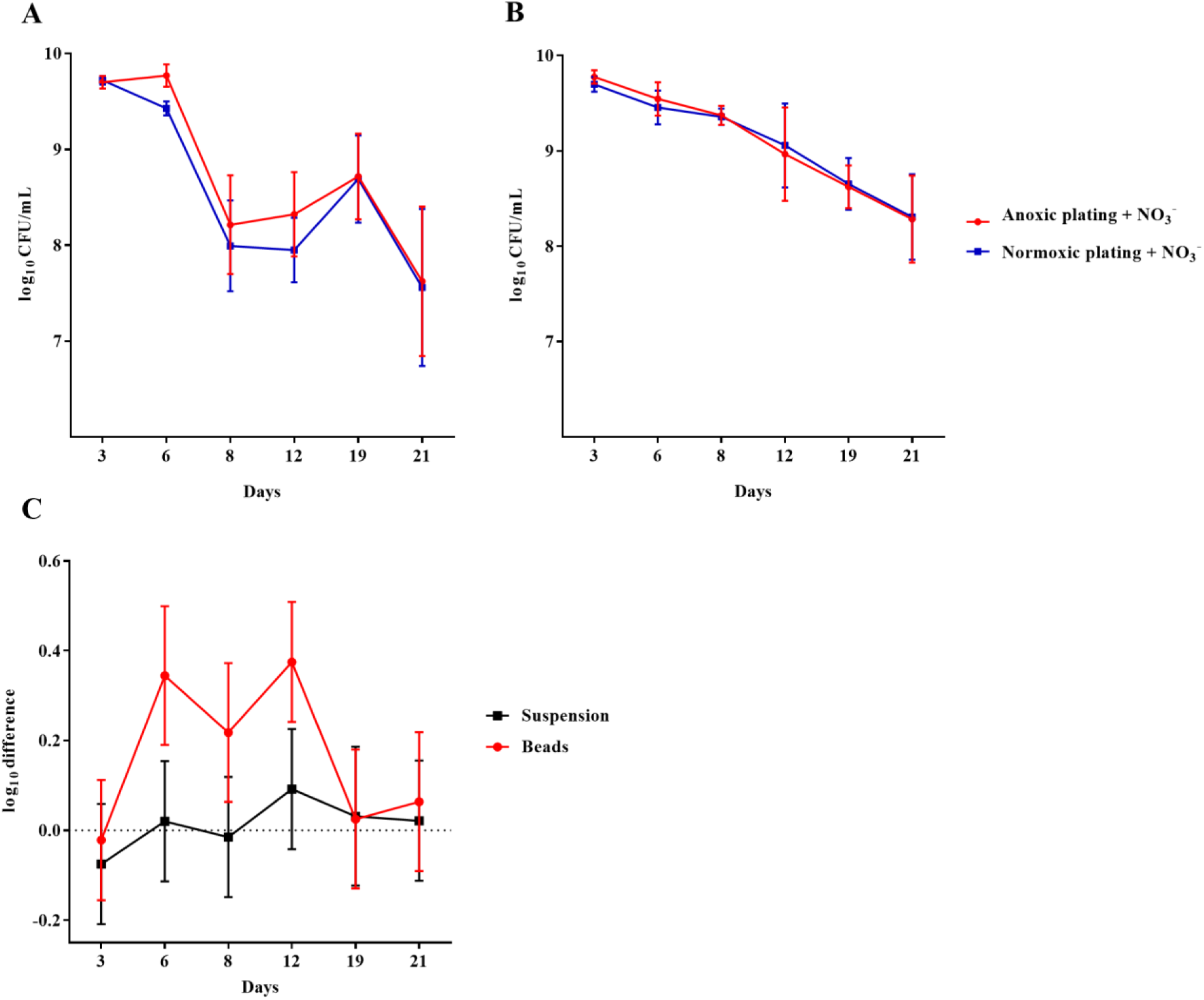
An oxygen intolerant subpopulation of *Pseudomonas aeruginosa* (PAO1) was generated in the bead biofilm model. Normoxic and anoxic colony forming units per milliliter (CFU/mL) of PAO1 over 21 days from the beads (A) and the surrounding suspension (B). Symbols with error bars indicate the mean ± SEM (n = 4). + NO_3_^-^ refers to the addition of 10 mM KNO_3_ to LB agar plates. The log difference (C) represents the difference in mean log CFU/mL between plating methods from the suspension and the beads, respectively. The log difference was significantly higher (p = 0.003, linear regression) in the beads than the surrounding suspension. Symbols with error bars indicate the mean + confidence intervals.

*P. aeruginosa* was also grown anoxically or normoxically on a filter biofilm model using LB plates supplemented with 10 mM NO_3_^-^ over a period of 17 days. CFU/mL was then determined using anoxic and normoxic plating, as above (Fig. 3A and 3B, respectively). The log difference between plating methods was significantly (p = 0.01) higher for anoxically conditioned biofilms than normoxically conditioned biofilms (Fig. 3C). The fraction of DTC *P. aeruginosa* ranged from six to 23 % from day three to 17 (Fig. S1).

**Fig. 3.**
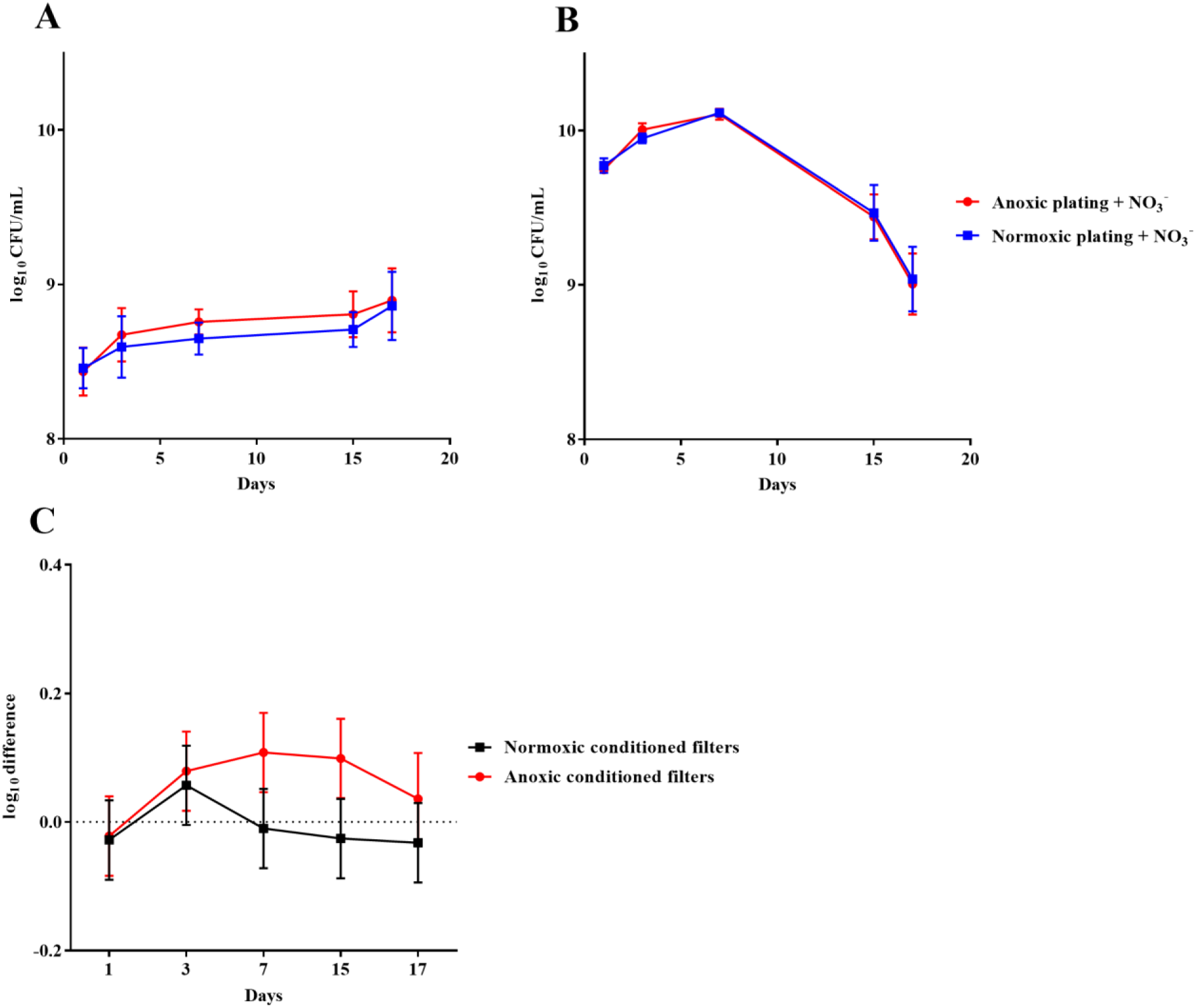
An oxygen intolerant subpopulation of *Pseudomonas aeruginosa* (PAO1) was generated in the filter biofilm model. Normoxic and anoxic colony forming units per milliliter (CFU/mL) of PAO1 over 17 days from anoxically (A) and normoxically (B) conditioned filters. Symbols with error bars indicate the mean ± SEM (n = 4). + NO_3_^-^ refers to the addition of 10 mM KNO_3_ to LB agar plates. The log difference (C) represents the difference in mean log CFU/mL between plating methods from the normoxically and anoxically conditioned filters, respectively. The log difference was significantly higher (p = 0.01, linear regression) in the anoxically conditioned filters than the normoxically conditioned. Symbols with error bars indicate the mean + confidence intervals.

Similar results were obtained for anoxically and normoxically conditioned planktonic batch cultures of *P. aeruginosa* over a period of 28 days (Fig. 4A and 4B, respectively). The log difference between plating methods was significantly (p < 0.0001) higher for anoxically conditioned batch cultures (Fig. 3C). The fraction of DTC bacteria ranged from 60 to 90 % from day nine to 21 (Fig. S1).

**Fig. 4.**
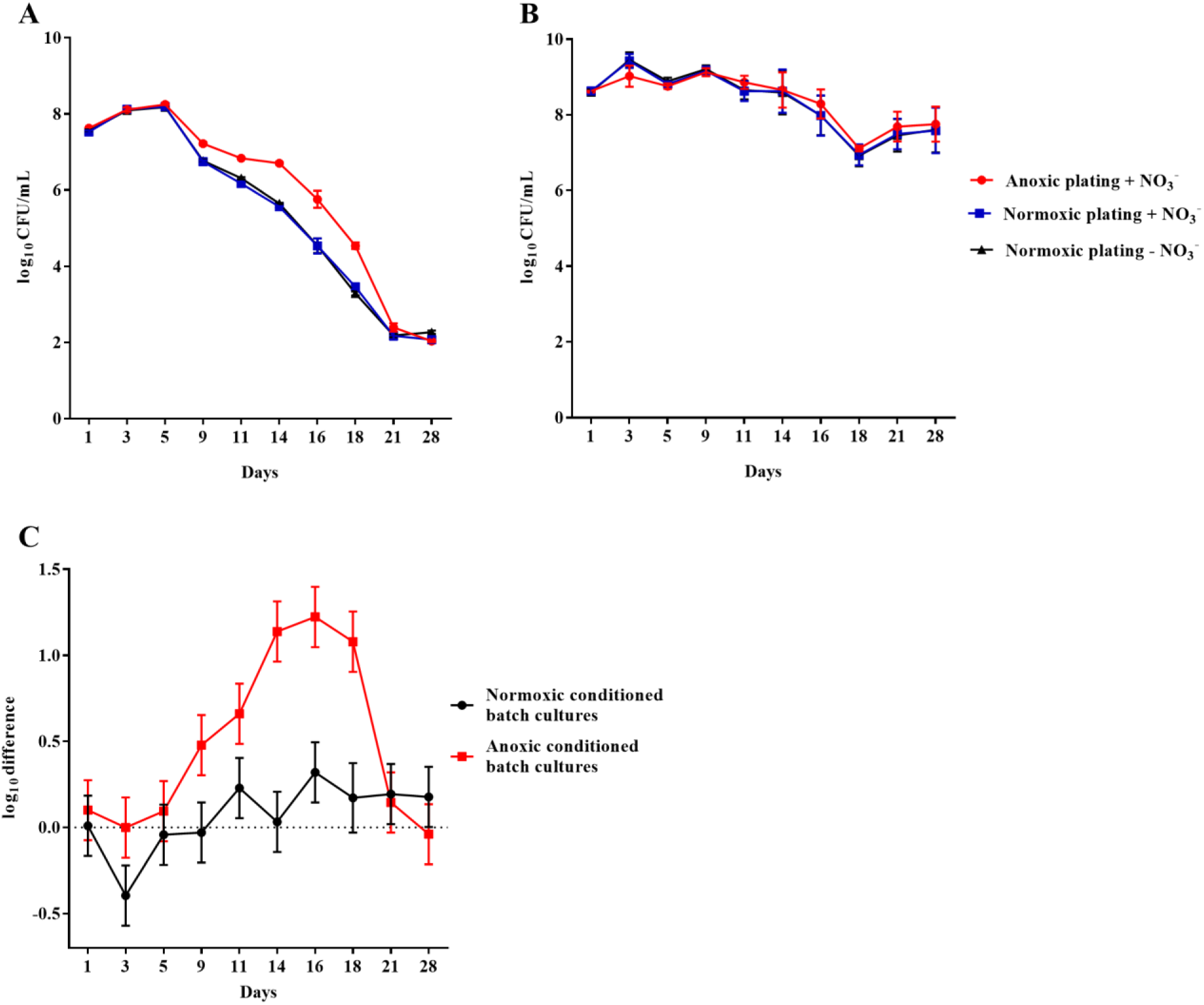
An oxygen intolerant subpopulation of Pseudomonas aeruginosa (PAO1) was generated in the planktonic batch cultures. Normoxic and anoxic colony forming units per milliliter (CFU/mL) of PAO1 over 28 days from anoxically (A) and normoxically (B) conditioned batch cultures. Symbols with error bars indicate the mean ± SEM (n = 4). ± NO_3_^-^ refers to the addition of 10 mM KNO_3_ to LB agar plates. The log difference (C) represents the difference in mean log CFU/mL between plating methods from the normoxically and anoxically conditioned batch cultures, respectively. The log difference was significantly higher (p < 0.0001, linear regression) in the anoxically conditioned batch cultures than the normoxically conditioned. Symbols with error bars indicate the mean + confidence intervals.

The effect of anoxic conditioning was further investigated on colonies of *P. aeruginosa* grown on LB plates supplemented with 10 mM NO_3_^-^ under anoxic and normoxic conditions over a period of 20 days (Fig. 5A and 5B, respectively). The log difference between plating methods was significantly higher (p < 0.0001) for anoxically conditioned colonies (Fig. 5C). The fraction of DTC *P. aeruginosa* ranged from 12 to 46 % from day six to 20 (Fig. S1).

**Fig. 5.**
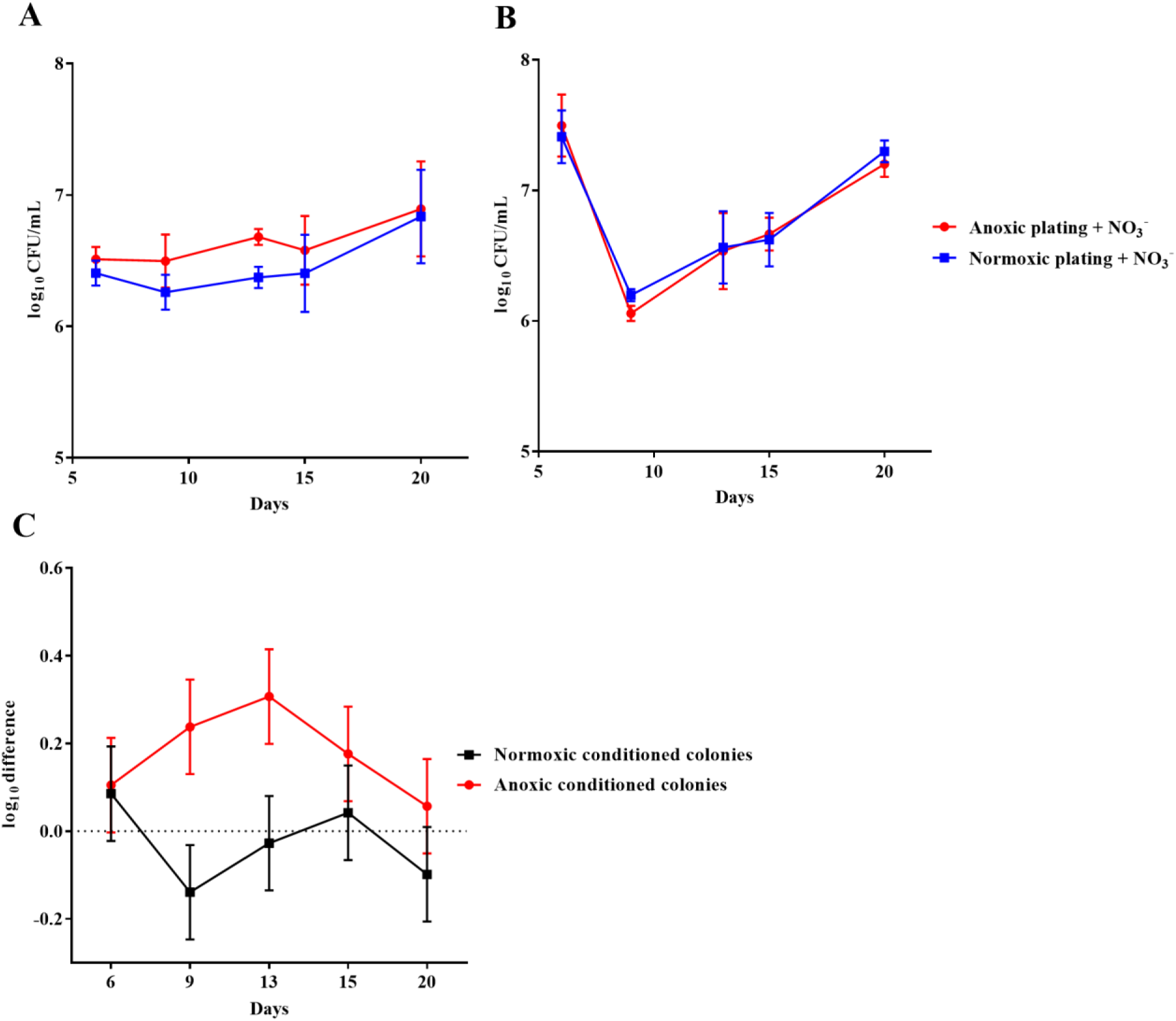
An oxygen intolerant subpopulation of *Pseudomonas aeruginosa* (PAO1) was generated in colonies. Normoxic and anoxic colony forming units per milliliter (CFU/mL) of PAO1 over 20 days from anoxically (A) and normoxically (B) conditioned colonies. Symbols with error bars indicate the mean ± SEM (n = 3). +NO_3_^-^ refers to the addition of 10 mM KNO_3_ to LB agar plates. The log difference (C) represents the difference in mean log CFU/mL between plating methods from the normoxically and anoxically conditioned colonies, respectively. The log difference was significantly higher (p < 0.0001, linear regression) in the anoxically conditioned colonies than the normoxically conditioned. Symbols with error bars indicate the mean + confidence intervals.

### 10 mM NO_3_^-^ in LB plates did not affect CFU/mL in batch cultures of *P. aeruginosa*

To ensure that presence of NO_3_^-^ did not affect the number of CFU generated on LB plates, CFU/mL was determined anoxically (LB plates + 10 mM NO_3_^-^) and normoxically (LB plates ± 10 mM NO_3_^-^) from 24-hour-old batch cultures of *P. aeruginosa*. No difference was observed between type of plating (p = 0.93): anoxic log CFU/mL + NO_3_^-^ = 9.96 (± 0.22 SD), normoxic log CFU/mL + NO_3_^-^ = 9.91 (± 0.10 SD) and normoxic log CFU/mL - NO_3_^-^ = 9.91 (± 0.14 SD). Furthermore, there was no effect of NO_3_^-^ supplementation on normoxic plating in the prolonged experiment with 28-day-old batch cultures (Fig. 4A and 4B). It was not possible to detect growth of *P. aeruginosa* on LB plates under anoxic conditions without NO_3_^-^, but growth was observed when these plates subsequently were placed under normoxic conditions.

### Oxidative stress restricted the growth of anoxically conditioned *P. aeruginosa* when re-grown in a normoxic environment

Bacteria from 16-day-old anoxically and normoxically conditioned batch cultures of *P. aeruginosa* were plated on 10 mM NO_3_^-^ LB plates ± sodium pyruvate and ± catalase to investigate whether formation of ROS contributed to the DTC state induced by anoxic conditioning. Presence of sodium pyruvate as a ROS scavenger (Mizunoe et al., 1999) significantly (p = 0.04) increased normoxic CFU/mL compared to plating without sodium pyruvate (Fig. 6B). This was not the case in 24-hour-old batch cultures of *P. aeruginosa* (p = 0.62), indicating that the effect was restricted to anoxically conditioned cells (Fig. 6A). Presence of catalase also significantly (p = 0.03) improved normoxic plate counts compared to normoxic plating without catalase (Fig. 6B). These results show that hydrogen peroxide is involved in the induction of a DTC state created by anoxic conditioning. To add proof to this hypothesis, a similar experiment was carried out with a catalase deficient *P. aeruginosa* mutant (Δ*katA* PAO1), which is more susceptible to oxidative stress (Hassett et al., 1999;Jensen et al., 2014). The log difference between plating methods for the Δ*katA* mutant and reference strain was significantly (p = 0.00002, unpaired t-test) different. These log difference values were 3.02 ± 0.05 SD and 1.61 ± 0.07 SD for Δ*katA* PAO1 and the reference strain, respectively (Fig. 6C). Moreover, measurement of ROS and optical density (OD) was carried out for *P. aeruginosa* at day 1, 8 and 16 from anoxically and normoxically conditioned batch cultures. These results showed creation of ROS during normoxic growth. The longer PAO1 was anoxically conditioned; the lower was the following normoxic growth, as measured by OD. This effect was less pronounced for normoxically-conditioned cells (Fig.7 A-C).

**Fig. 6.**
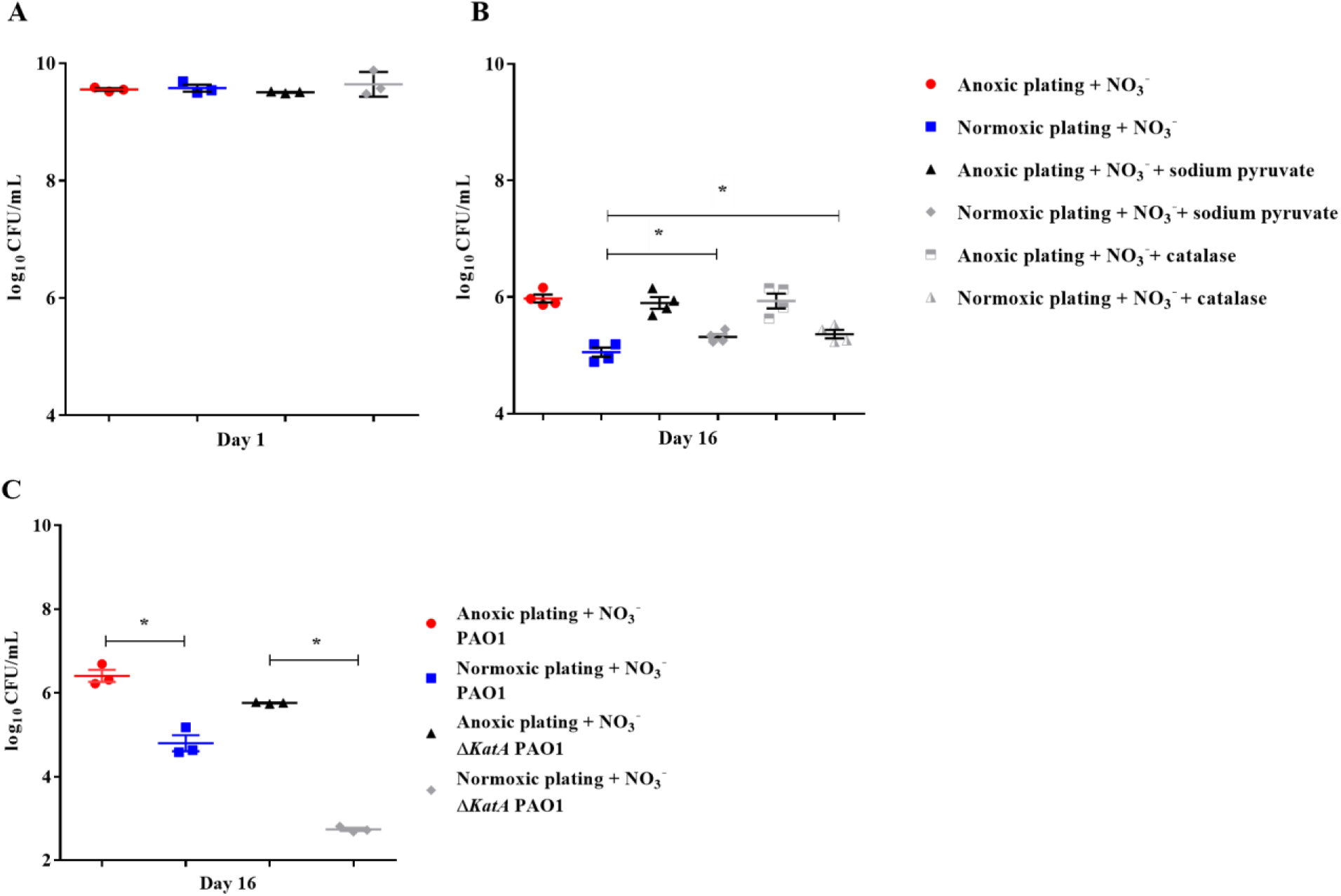
Reactive oxygen species generates a DTC subpopulation of *Pseudomonas aeruginosa* (PAO1) Normoxic and anoxic colony forming units per milliliter (CFU/mL) were determined for 24-hour-old batch cultures (A) and 16-day-old anoxically conditioned batch cultures (B) of PAO1. Normoxic CFU/mL was determined ± 0.3% sodium pyruvate (A and B), while addition of catalase was only tested for 16-day-old batch cultures. Symbols with error bars indicate the mean ± SEM (n = 4). +NO_3_^-^ refers to the addition of 10 mM KNO_3_ to LB agar plates. There was a significant difference between normoxic plating ± 0.3 % sodium pyruvate (p = 0.04, one-way ANOVA test) and normoxic plating ± catalase (p = 0.03, one-way ANOVA test). Normoxic and anoxic determination of CFU/mL were determined for 16-day-old anoxically conditioned batch cultures of PAO1 and *ΔkatA* PAO1 (C). Symbols with error bars indicate the mean ± SEM (n = 3). Significant difference between anoxic and normoxic plating with both PAO1 and *ΔkatA* PAO1 (p = 0.0014 and p < 0.0001, respectively, one-way ANOVA test).

**Fig. 7.**
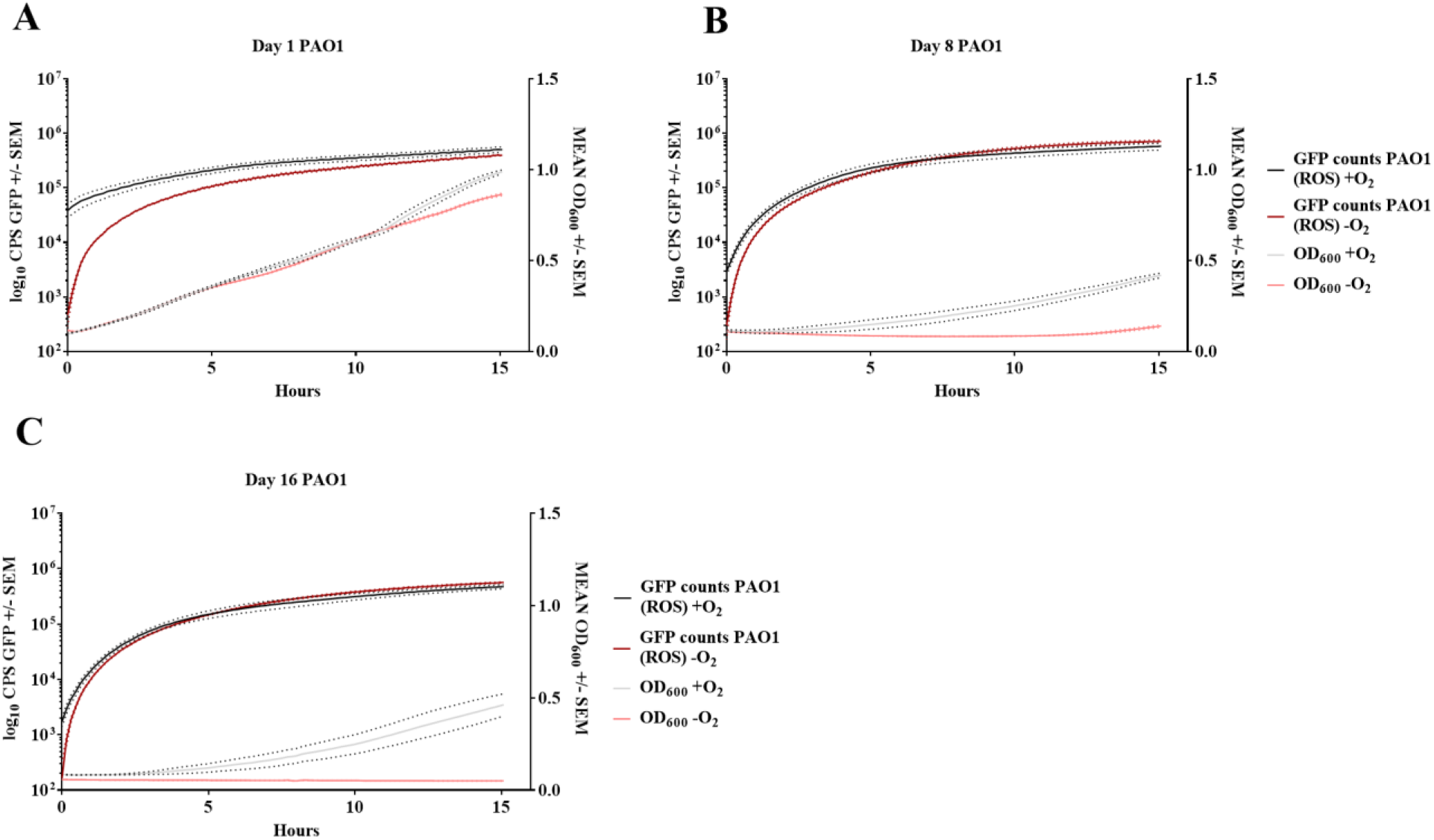
Increase in reactive oxygen species affects growth of anoxically conditioned *Pseudomonas aeruginosa* (PAO1) over time. Determination of reactive oxygen species (ROS) and optical density (OD) at day 1 (A), 8 (B) and 16 (C) for anoxically (-O_2_) and normoxically (+O_2_) conditioned PAO1. ROS was measured as counts per second (CPS) when 2′,7′dichlorodihydrofluorescein diacetate was converted to 2′7′-dichlorofluorescein. Growth was measured with optical density (OD_600_) simultaneously to ROS measurement. Data is generated from 99 continuously measurements presented as a solid line with dots representing ± SEM (n = 4).

### Anoxic conditioning affects *Staphylococcus aureus* and *Staphylococcus epidermidis*, but not *Escherichia coli* and *Enterococcus faecalis*

The effect of anoxic conditioning was then tested on a selected group of pathogens to determine if this phenomenon was restricted to *P. aeruginosa. S. aureus* (methicillin susceptible) was tested as described in the *P. aeruginosa* filter biofilm setup. Plate counts for anoxically and normoxically conditioned filters was determined (Fig. 8A and 8B, respectively). The log difference between plating method was significantly higher (p < 0.001) when plate counts were performed from anoxically conditioned filter biofilms than from normoxically conditioned filter biofilms (Fig. 8C). The fraction of DTC *S. aureus* ranged from three to 89 % from day one to day 17 (Fig. S1). Since both *P. aeruginosa* and *S. aureus* demonstrated effects of anoxic conditioning, a smaller experiment was initiated to test other pathogens. The effect was determined on LB plates supplemented with 10 mM NO_3_^-^ ± sodium pyruvate to investigate if ROS were involved in the lack of growth. Both *S. aureus* (MRSA) and *S. epidermidis* showed effects of anoxic conditioning, resulting in an significant (p = 0.01 and p = 0.03, respectively) increase in CFU/mL during anoxic plating compared to normoxic plating (Fig. S2). The fraction of DTC *S. aureus* (MRSA) and *S. epidermidis* was 36 and 42 % at day 9, respectively. There was no observed effect of sodium pyruvate during normoxic plating for these two strains. In the case of *E. coli* and *E. faecalis*, there was no observed effect of anoxic conditioning (Fig. S3).

**Fig. 8.**
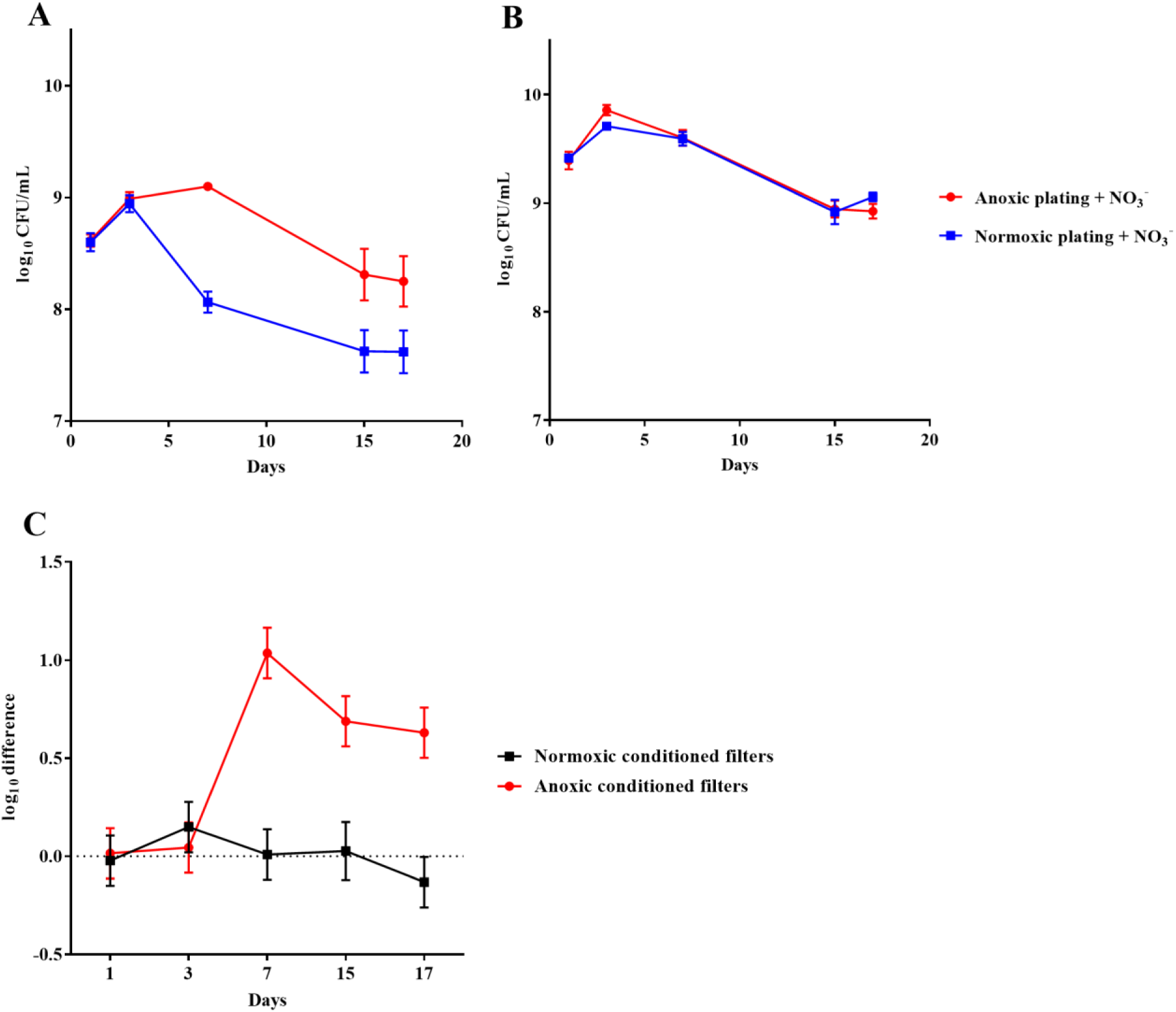
An oxygen intolerant subpopulation of *Staphylococcus aureus* (methicillin susceptible) was generated in the filter biofilm model. Normoxic and anoxic colony forming units per milliliter (CFU/mL) of *Staphylococcus aureus* over 17 days from anoxically (A) and normoxically (B) conditioned filters. Symbols with error bars indicate the mean ± SEM (n = 4). +NO_3_^-^ refers to the addition of 10 mM KNO_3_ to LB agar plates. The log difference (C) represents the difference in mean log CFU/mL between plating methods from the normoxically and anoxically conditioned filters, respectively. The log difference was significantly higher (p < 0.001, linear regression) in the anoxically conditioned filters than the normoxically conditioned. Symbols with error bars indicate the mean + confidence intervals.

## Discussion

### DTC and VBNC *P. aeruginosa*

In the current study, we demonstrate that plate counting with *P. aeruginosa* is significantly affected by anoxic conditioning when cultured in the presence of atmospheric oxygen levels. A DTC state was created after only a few days of anoxic conditioning in both biofilm and planktonic models. DTC *P. aeruginosa* was observed in the beads from the bead biofilm model, but not in the surrounding suspension, despite that the cultures had full access to atmospheric oxygen. This indicates that oxygen restriction within a biofilm generates a DTC subpopulation. The DTC state, detected as increased growth during anoxic plating with NO_3_^-^, has previously been generated in planktonic *P. aeruginosa* by energy starvation after cultivation without O_2_ as electron acceptor for aerobic respiration (Binnerup, 1993). *In vitro* biofilms of *P. aeruginosa* may contain internal anoxic zones (Walters et al., 2003;Kolpen et al., 2016), so we hypothesized that anoxia inside the biofilm contributes to the induction of DTC *P. aeruginosa*. Accordingly, DTC *P. aeruginosa* was only observed in the filter biofilm model and the colony model when the model was anoxically conditioned. Interestingly, we found that 99.68 % of the population was VBNC with the LIVE/DEAD assay after anoxic conditioning. LIVE/DEAD staining is based upon membrane permeability and is only an approximation of true viability. Cells that were stained both red and green (resulting in yellow) were counted as dead cells but may still be viable.

Approximately 50 % of the population in the alginate beads was DTC even though they were kept in normoxically-conditioned vials. In comparison, the fraction of DTC cells was approximately 90 % in the anoxically conditioned batch cultures. The difference (90 % vs. 50 %) may be explained by the fact that a majority of the bacteria in the beads are peripherally located where they have increased access to oxygen compared to the center of the beads (Sønderholm et al., 2017). In comparison, bacteria from the anoxically conditioned batch cultures were fully deprived of O_2_. In the present study, the effect of anoxic conditioning was restricted to an intermediate period. This is probably due to the static methodological setup. Dynamic processes, such as entry of nutrients and removal of waste, occur during infection and are not modeled here (Brown et al., 2008). Nevertheless, this is not the first time a “resuscitation window” has been described for non-culturable bacteria. Similar studies have shown that VBNC bacteria can only be detected in an intermediate period of culturing (Pinto et al., 2015).

### ROS restricts growth of anoxically conditioned *P. aeruginosa*

The DTC fraction of bacteria in this study was O_2_ intolerant given that growth only was achievable when NO_3_^-^ served as an alternative electron acceptor during anaerobic respiration. This led us to investigate whether this phenomena was due to oxidizing properties of ROS created by incomplete reduction of O_2_ during aerobic respiration (Fenchel and Finlay, 2008). Sodium pyruvate increased counts for anoxically conditioned *P. aeruginosa* during normoxic plating (roughly 43 % of the DTC population) and we believe that it was due to its properties as a ROS scavenger (Mizunoe et al., 1999). Accordingly, 0.3 % sodium pyruvate has been used to resuscitate non-culturable populations of *S. aureus* (Pasquaroli et al., 2013). Catalase also increased counts during normoxic plating (roughly 50 % of the DTC population), indicating that hydrogen peroxide is involved in the induction of a DTC state. The effect of ROS was further confirmed by the reduced growth of a catalase A deficient *P. aeruginosa* mutant (Δ*katA* PAO1) after anoxic conditioning. Finally, measurements of ROS along with OD showed that anoxically conditioned batch cultures of *P. aeruginosa* created more ROS when regrown under atmospheric conditions the longer they were deprived from oxygen, resulting in an accumulation of ROS and a reduction in growth represented by lower OD measurements. These results indicate that accumulation of ROS during aerobic respiration has an impact on normoxic plate counting when *P. aeruginosa* has been anoxically conditioned. Furthermore, it seems that hydrogen peroxide is involved in this loss of culturing. It has been suggested that VBNC cells cannot be resuscitated by addition of ROS scavengers and that “revived” cells in the presence of ROS scavengers are only injured cells and not VBNC cells (Pinto et al., 2015). It was not possible to determine whether DTC bacteria in this study were in an injured state.

### *S. epidermidis* and *S. aureus* also become DTC after anoxic conditioning

Additional experiments were conducted to elucidate whether anoxic conditioning generated DTC sub-populations in other facultative pathogens. Filter biofilms of methicillin susceptible *S. aureus* showed the same effects of anoxic conditioning as *P. aeruginosa*. These experiments were also repeated for *S. epidermidis, S. aureus* (MRSA), *E. coli* and *E. faecalis*. Both *S. epidermidis* and *S. aureus* were significantly (p = 0.01 and p = 0.03, respectively) affected by anoxic conditioning, whereas *E. coli* and *E. faecalis* were not. The results in this study points toward an intermediate period where anoxic conditioned bacteria are benefitted by anoxic incubation with nitrate as alternative electron acceptor. We cannot rule out that we missed the period where *E. coli* and *E. faecalis* were DTC, given that they were tested at selected days. No effect of sodium pyruvate was found during normoxic plating for any of these tested organisms, suggesting that ROS is not involved in the loss of culturability for these pathogens, but additional ROS scavengers should be tested.

## Conclusion

A non-culturable state can be induced by several physiological stresses and our knowledge in this area is expanding, though far from fully resolved. In this study, we demonstrate that anoxic conditioning generates a VBNC subpopulation in *P. aeruginosa* batch-cultures. Furthermore, we demonstrate that anoxic conditioning generates DTC sub-populations of *P. aeruginosa, S. aureus* and *S. epidermidis* during biofilm growth. The DTC population was only able to grow under anoxic conditions in the presence of NO_3_^-^ as an alternative electron acceptor. In the case of *P. aeruginosa*, this phenomenon was explained by creation of lethal amounts of ROS during aerobic respiration and the bacteria’s inability to neutralize it.

## Data availability

The data generated in the current study are available from the corresponding author on reasonable request.

## Author contributions

LK performed the majority of the experiments. MK, MS, BFG, SC and KNK also performed experiments. TB, PØJ, KNK AK, and LK conceived and designed experiments. LK wrote the manuscript. MA and PØJ performed statistics. All authors analyzed data. All authors contributed to and corrected the manuscript.

