## Supplemental Figure S1 for "Oxygen Restriction Generates Difficult-To-Culture Pathogens"

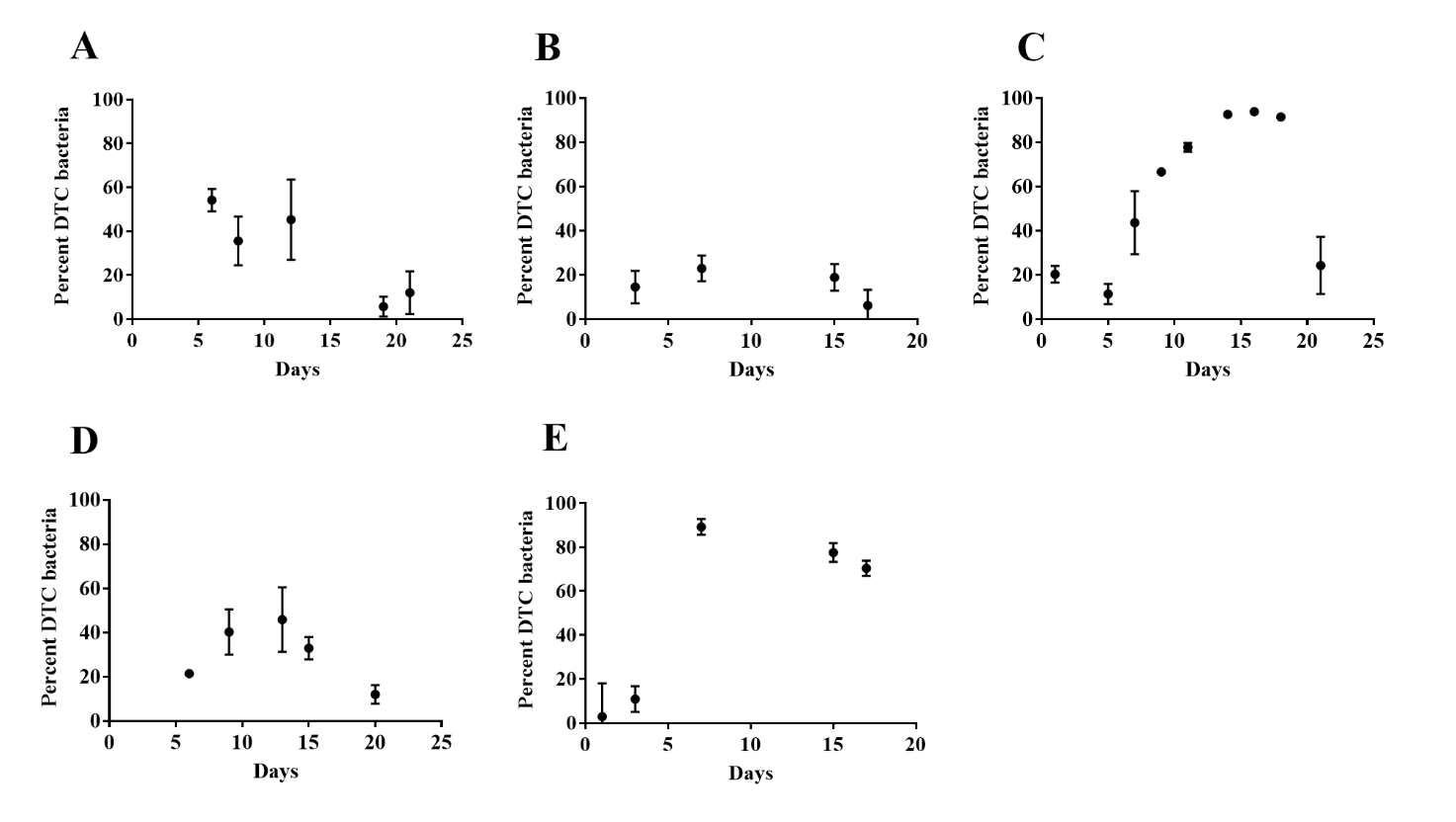

Fig. S1 – Percentage distribution of DTC bacteria over time in different growth models

The percentage of DTC *Pseudomonas aeruginosa* after anoxic conditioning in the bead biofilm model (A), filter biofilm model (B), batch cultures (C), colonies (D). The percentage of DTC *Staphylococcus aureus* after anoxic conditioning in the filter biofilm model (E).
