## Supplemental Figure S2 for "Oxygen Restriction Generates Difficult-To-Culture Pathogens"

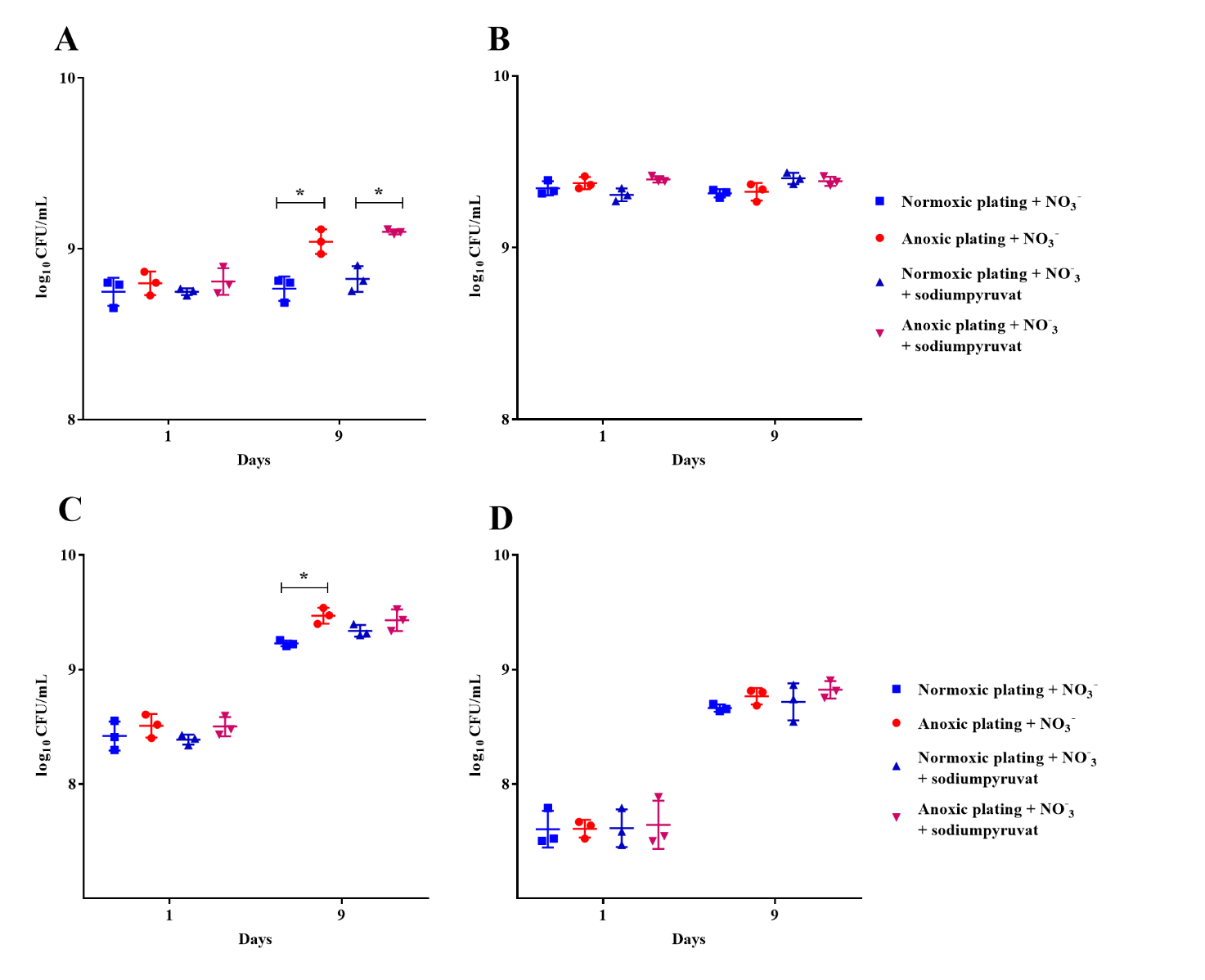

S2 - An oxygen intolerant subpopulation of S*taphylococcus aureus* (MRSA) and *Staphylococcus epidermidis* was generated in the filter biofilm model.

Normoxic and anoxic (± sodium pyruvate) determination of CFU/mL were determined for 1 and 9-day-old anoxically (A) and normoxically (B) conditioned filters with *Staphylococcus aureus* (MRSA) and anoxically (C) and normoxically (D) conditioned filters with *Staphylococcus epidermidis*. Significant difference between anoxic and normoxic plating ± sodium pyruvate at day 9 (p = 0.01 and p < 0.01, respectively, one-way ANOVA test) for *S. aureus*. Significant difference between anoxic and normoxic plating at day 9 (p = 0.003, one-way ANOVA test) for *S. epidermidis*. Symbols with error bars indicate the mean ± SEM (n = 3). +NO_3_^-^ refers to the addition of 10 mM KNO_3_ to LB agar plates.
