## Supplemental Figure S3 for "Oxygen Restriction Generates Difficult-To-Culture Pathogens"

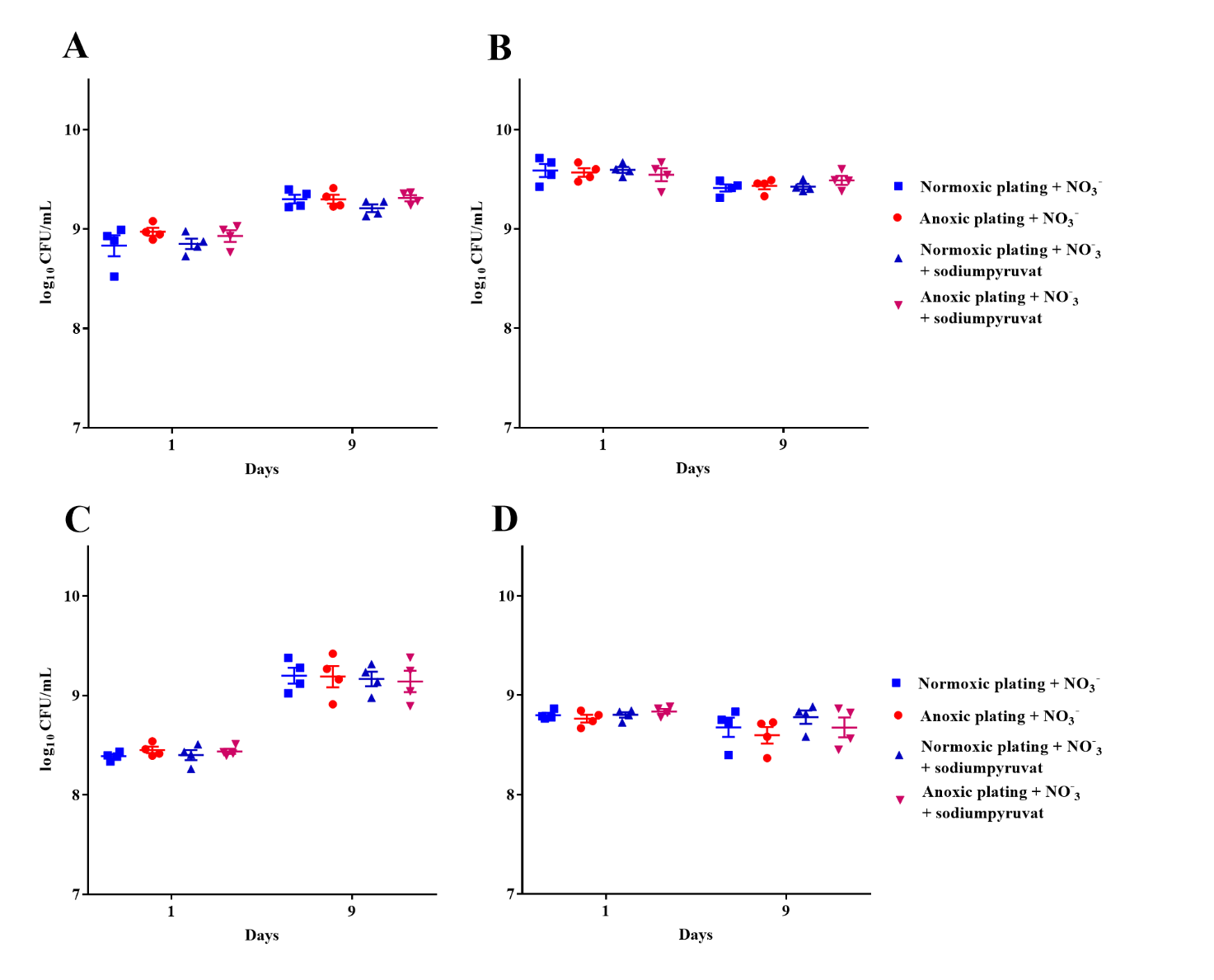

S3 – Anoxic conditioning did not affect E*scherichia coli* or *Enterococcus faecalis* during growth in the filter biofilm model.

Normoxic and anoxic (± sodium pyruvate) determination of CFU/mL were determined for 1- and 9-day-old anoxically (A) and normoxically (B) conditioned filters with *E. coli* and anoxically (C) and normoxically (D) conditioned filters with *E. faecalis*. No difference between types of plating (one-way ANOVA test). Symbols with error bars indicate the mean ± SEM (n = 3). +NO_3_^-^ refers to the addition of 10 mM KNO_3_ to LB agar plates.
